# Differentiable Vertex Model: Exploring Gradient-Based Optimization for Tissue Morphogenesis

**DOI:** 10.64898/2026.05.07.723189

**Authors:** Lars Erik J. Skjegstad, Steven Oud, Renske M. A. Vroomans, Julius B. Kirkegaard

## Abstract

Vertex models are widely used within the field of developmental biology to study tissue morphogenesis. These models are well-suited for modeling deformation at the cellular level where movement is driven by local forces. However, understanding how these microscopic movements coordinate to yield macroscopic phenomena such as the shapes of entire tissues remains a challenge. Here we study a top-down approach using differentiable programming on a simplified vertex model of a laminar tissue, and investigate whether the attributes of individual cells can be tuned to make the mesh as a whole acquire a predefined shape. We let the mesh evolve according to simple rules defined by the input to each polygon, and evaluate the resulting shape against a target boundary. Additionally, we show how the high degeneracy of the output can be reduced by constraining the polygon distributions: first, by adding simple penalties on tissue-wide attributes; and second, by dividing the tissue into regions, within which we bias the attributes toward characteristic values. Our study shows how a simple vertex model can be combined with differentiable programming to model developing tissues, and provides insight into the way individual cells must coordinate to yield macroscopic phenomena such as pre-programmed shapes.

## INTRODUCTION

Vertex models are commonly used within the field of developmental biology for modeling cells within tissues, particularly tightly-packed, confluent tissues such as animal epithelium and plant epidermis [1–3]. They are among the simplest mechanical cell model formalisms to implement, which makes it easy to test different hypotheses about the role of various mechanical forces during morphogenesis [3]. In vertex models, cell configurations are described by the positions and connections of vertices, and tissues can thus be represented as meshes where each edge represents a cell membrane or cell wall, typically modeled as an elastic or viscoelastic spring connecting two vertices (point masses). The positions of the vertices are updated based on the forces acting on them, such as internal, osmotic pressure (turgor in plants), and membrane or cell wall tension. These models can also incorporate more complex phenomena, such as cell intercalation and cell division [4]. One of the areas in which vertex models have seen widespread use is that of *morphogenesis*, the development of shape in biological systems [1, 2]. Morphogenesis is typically simulated by integrating the forces acting on all vertices forward in time, with the tissue as a whole changing based on their microscopic interactions at the vertices [2, 3]. When modeling plant tissues, cells must be prevented from sliding past each other by forcing neighboring cells to share edges. Plant cells also do not shrink through active contraction, which can be ensured by not allowing the rest lengths of the edges to decrease. Simple vertex models have, for example, been used to test the effect of cell-division rules on the heterogeneity of tissue growth [5], the effect of different cell types on root bending [6], and the role of cell-autonomous and non-autonomous processes in lateral organ outgrowth [7]. Recent studies have integrated vertex models with additional processes: hormone transport dynamics to explain the patterning of veins in growing leaves [8, 9] and water fluxes to explain the outgrowth of plant organs from the stem cell zone [10].

Morphogenetic processes are complex from a modeling perspective, given that many biological as well as physical mechanisms can be included in a model. The number of variables and parameters can potentially be vast, given that multiple parameters (such as edge elasticity and area growth rate) can be assigned to each individual cell, resulting in significant computational cost, as well as increased difficulty in matching models to data. Multiscale approaches [11–13] seek to explicitly couple subcellular, cellular, and tissue-level dynamics within a unified framework but can suffer from the *tyranny of scales* problem [14, 15]: Forces are strongly coupled, both locally (neighboring cells pushing against each other) and across scales (local cell shape changes due to, e.g., cell wall bending, leading to entire tissue regions expanding differently) [16]. These phenomena make parameter tuning very challenging and obfuscate model interpretability. Lastly, macroscopic forces are hard to control, which can lead to, e.g., high compression of cells near rigid surfaces (such as points of attachment of the modeled tissue to a stem).

One ongoing topic of research on modeling developmental biology is the link between interactions at the cellular level and the emergent *shapes* of mature tissues [17–20]. One constraint that forward approaches share is that they require some knowledge of, or hypothesis about, the microscopic forces and growth patterns in the tissue, but if only the final shape of the tissue is known, inferring these becomes difficult. To enhance our understanding of the morphogenesis of shapes, we instead study an alternative approach to the problem. In this paper, we employ a simplified model in which smaller-scale phenomena (such as stress within fibers, cell wall mechanics, and microtubule orientation) are abstracted away [16]. We thus always assume that the forces involved at these orders of magnitude are at equilibrium, motivated by the fact that the time scale at which these phenomena occur is much smaller than the time scale of growth at the tissue level (i.e., tissue growth is a quasi-static process). As a result, cell morphogenesis as modeled in this study is implicitly coordinated across the entire tissue. Notably, our approach is *top-down*: We let the shape of the mature tissue dictate the behavior of the individual cells, and from the resulting configuration of the cells infer knowledge on their individual characteristics, such as size and shape anisotropy. The model requires only a few parameters per cell, which act as aggregate lower-level effects. To accomplish this, we let the cell characteristics constitute the trainable parameters, and the final shape constitute the target for model accuracy, using differentiable programming to evolve the meshes. This approach has previously been used by [21] to study the relationship between local and macroscopic effects in biological tissues using adhesive soft spheres. Here we use a vertex model to explore this relationship.

Practically, we propose a dual use of differentiable programming: (1) Instead of formulating the vertex model explicitly in terms of forces, we specify the system in terms of potentials, the derivatives of which determine the physical evolution of the system, and (2) we select final morphogenetic metrics for which we wish to optimize, aiming to achieve this by calculating the gradients of these metrics through the entire temporal evolution of the simulation.

Fig. 1 illustrates the model framework. We define a forward model as a vertex model where the mesh evolves as a function of the learnable input parameters, which in our model are the area (shown in green) and anisotropy (shown in orange) for each individual polygon. The final shape of the mesh is then compared to a predetermined target boundary, e.g., the shape of a flower petal (both indicated with dotted lines). A loss value is computed based on the discrepancy between the shapes, and gradients with respect to the loss value are computed by automatic differentiation in JAX [22]. Finally, the gradients are used to update the parameters for each polygon. The process is repeated until convergence.

**FIG. 1:**
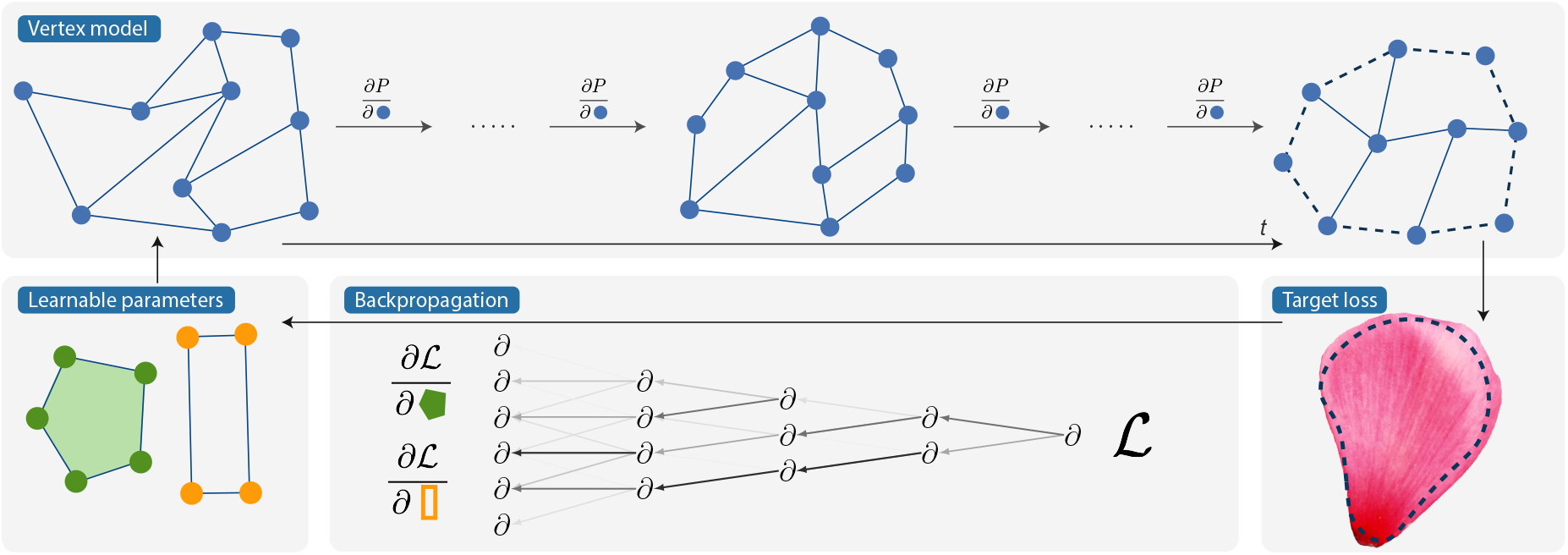
Illustration of the model framework. The mesh evolves via vertex-position derivatives of a potential function parameterized by learnable variables. The resulting shape is compared to that of a predefined target (e.g., a flower petal). Gradients of the loss with respect to the learnable parameters (e.g., areas and anisotropies) are then used to update the parameters. All derivatives are computed using automatic differentiation through the computational graph.

Our model is a simplification of the morphogenetic process, not only due to the number of parameters being low relative to that of microscopically specified models, but also in that the number of polygons is both constant and considerably lower than the number of cells in a mature tissue (which contains millions of cells). This is partly done because remeshing is inherently non-differentiable, and as such, differentiable programming is not well suited for modeling discrete events such as cell division or extrusion. Although differentiable programming is applicable to fixed cell division programs in principle, we note that the problem of cell divisions would require a discrete optimization approach, and we focus this study on pure differentiable programming.

## MODEL

The tissue is represented as a 2D apical vertex model [1], created by initializing points randomly with a given density parameter. A Voronoi tessellation is computed from the points, and subsequently clipped by a semicircle (see Fig. 2a). We let the line generated by the straight edge constitute the tissue base, i.e., the boundary which separates the tissue from the rest of the organism, to which the tissue is attached. This procedure is used to generate initial meshes for all experiments, their shape being inspired by early-stage organ primordia emanating from the stem cell niche (meristem) in plants.

**FIG. 2:**
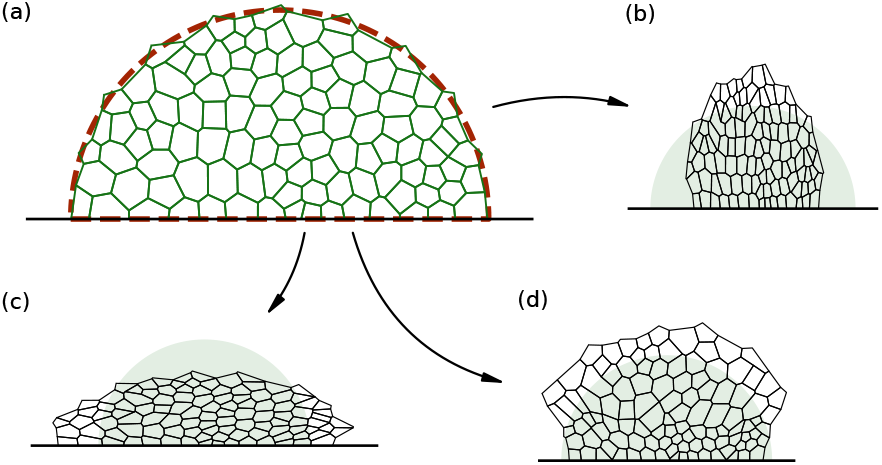
**(a)** An initial mesh. The dashed lines indicate the generating shape. **(b-d)** Output tissues from the `MORPH` program, using extreme goal metrics. The shape of the initial mesh is indicated in the background. **(b)** High, positive anisotropy. **(c)** High, negative anisotropy. **(d)** Top half has larger goal areas.

The system is defined by a (fixed-length) (*N*_*v*_, 2) array ***v*** of unique vertices, and each polygon ***p***_*i*_ is defined as an ordered list of indices into the array. The edges are implicitly defined by this ordering, shared between neighbors, and are constant, as we allow no topological changes in the mesh to occur. We thus do not model cell division, and let the polygons in our model represent *regions* of cells, where any change in the size and shape of a region should be interpreted as the combined effects of growth and division of the cells that the region represents. With these data structures, it is easy to perform computations on each individual polygon. Concretely, we consider the following metrics: the area *a*_*i*_ (computed using the shoelace formula) and the anisotropy

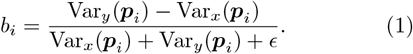

These metrics should be interpreted as emergent attributes resulting from lower-level interactions, and here serve as part of a naïve approach to describing the geometric properties of the polygons.

### Forward model

To simulate tissue deformation, we assign two input parameters to each polygon: a goal area *α*_*i*_ and a goal anisotropy *β*_*i*_. These represent the pre-programmed attributes that the polygons should attain after the morphing process (although the resulting values will also be constrained by the presence of neighboring polygons in the tissue). We consider simulations that take a fixed number of steps, *T*_m_, and at each step *t* we employ the simplest possible potentials by squared deviations,

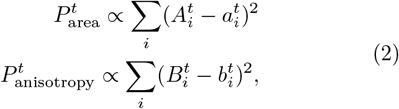

where 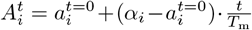 is a linear interpolation between the initial area and the goal area (*B*_*i*_ being defined analogously). This continuation scheme helps keep gradients bounded and stabilizes the simulation (reminiscent of gradient clipping). Thus, the parameters for all polygons, ***α*** and ***β***, determine the size and shape of the final mesh, whereas *T*_m_ controls the step size given the parameters. Additionally, the model necessitates a penalty on non-convex polygons. This term discourages edge crossings (although it does not guarantee that crossings are avoided) and is physically reminiscent of turgor pressure. For each polygon this potential is defined as a sum over its interior angles:

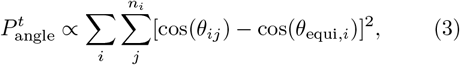

where *n*_*i*_ is the number of vertices in the *i*-th polygon, and 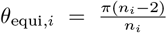 is the value of all angles in an equiangular polygon. At each step the vertex positions are updated by computing the gradients 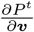 on the total potential

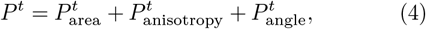

using intermediate adaptive time stepping to enhance simulation stability.

Importantly, the vertices at the base are fixed in *y* to emulate the expansion of the tissue in the upward direction. The vertices are, however, still free to move in *x*, which allows for lateral tissue movement.

We can now obtain morphed meshes by running the `MORPH` program on the initial mesh, defined by the initial vertices ***v***_init_:

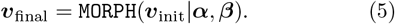

Figs. 2b-d demonstrate some examples of `MORPH` program output. In Fig. 2b, the goal anisotropy for all polygons is set to a high value, which results in highly elongated polygons, whereas in Fig. 2c, the goal anisotropies are set to a low value, resulting in flattened polygons. Fig. 2d shows the effects of assigning to the top-half polygons goal areas twice the values of the polygons in the lower half, resulting in a tissue where the upper polygons are considerably bigger than the lower ones. When manipulating the input parameters of each individual polygon, the tissue can be further tailored to take on a wide range of shapes. We now seek to do this by optimizing these parameters top-down for the mesh as a whole to match predetermined target shapes.

## RESULTS

### Shape optimization

Given a target boundary, we optimize the input goal parameters ***α*** and ***β***, and thus attempt to find a `MORPH` program that yields a mesh with a predetermined shape. We represent the boundary as a polygon ***t***, the shape of which is symmetric around the central vertical axis of the initial mesh. As we focus our attention on tissue morphogenesis rather than tissue growth, we use coordinates that scale with overall tissue expansion [23], and thus let the area contained by the target boundary be the same as that of the initial tissue. For each optimization step, the `MORPH` program is run once, and the loss is computed by comparing the boundary of the output mesh with the target boundary. The procedure results in an optimized mesh according to defined criteria.

The mapping from the initial to the final mesh is many-to-one, particularly due to the fact that only the boundary of the mesh is evaluated for the loss; indeed, it is easy to imagine local variations in the polygon shapes and sizes which together make up a mesh with exactly the same boundary. Due to the high non-convexity of the optimization problem, the model output parameters will often be correlated with the input parameters, and the choice of the initial values is thus non-trivial. Here, we consider the *Tutte embedding*, which we initially use to map the initial mesh into the target shape. This analytical mapping yields a distribution of polygons which only depends on the initial mesh and target boundary, with the guarantee that no edges cross. (Importantly, the mapping imposes no constraints given by the definition of the `MORPH` program.) After performing the embedding, we measure the individual areas and anisotropies directly, and use those as the initial guess in the shape optimization. (See Fig. S1 for an example comparison between the outputs of the embedding and the subsequent optimization). Although any choice of initial values will bias the output due to the aforementioned non-convexity of the problem, we note that such a choice will act as a restriction on the model output space, which in any case is necessary to make the model more biologically realistic. At a later stage we add more constraints motivated by the distributions of cells found in real tissues.

#### Optimization procedure

The optimization consists of the following main steps: First, the initial parameters ***α***^*n*=0^, ***β***^*n*=0^ are obtained from the Tutte embedding. Then, at each step *n*, the full `MORPH` program is run, and the vertices of the final tissue are obtained as

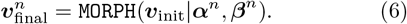

The loss is calculated by comparing the boundary vertices ***v***_b_ of the final tissue with the target boundary:

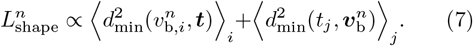

Here *d*_min_ is computed as the distance from a mesh (target) boundary vertex to the closest target (mesh) boundary edge. We assign an infinite loss to unphysical output, in particular, where two or more edges cross. Next, the gradients 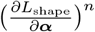,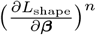 can be computed with respect to the loss, using automatic differentiation in JAX. Finally, the updated parameters ***α***^*n*+1^, ***β***^*n*+1^ are found by gradient descent. The optimizer uses the Adam algorithm with gradient clipping as well as a cosine decay learning rate schedule. See SI section III for details on calculations as well as parameter constraints.

We find that the model yields morphed tissues that match a range of different target boundaries. Fig. 3 shows four different examples: a tall isosceles trapezoid, a wide isosceles trapezoid, a petal-like shape, and a non-convex shape. Starting from the same initial mesh, the model converges on parameters that individually contribute to the overall shape of the output mesh such that the boundaries of the mesh and target largely overlap in all of these cases. The polygons are colored by the logarithmic deviations of the areas, 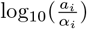 (see Fig. S3 for analogous plots for the anisotropies). Here we see that the goal metrics and the measured metrics differ. These residual discrepancies reflect constraints imposed by the `MORPH` program, and arise partly from the fact that not all target configurations are attainable from a given initial mesh. For instance, the morphed meshes favor convex polygons, leading to a natural limit on the space of shapes they can take on. (For an elaboration on model limitations, see SI section IV.) Neighboring polygons are also mechanically coupled, so changing the goal area or anisotropy of one polygon necessarily affects its neighbors. Some combinations of target shape and polygon-level metrics may therefore be mutually incompatible, leaving residual differences of physical origin between the imposed goals and the measured metrics. Finally, since the shape optimization problem is highly non-convex, some residual discrepancy may also arise from the local optimizer converging to a suboptimal `MORPH` program.

**FIG. 3:**
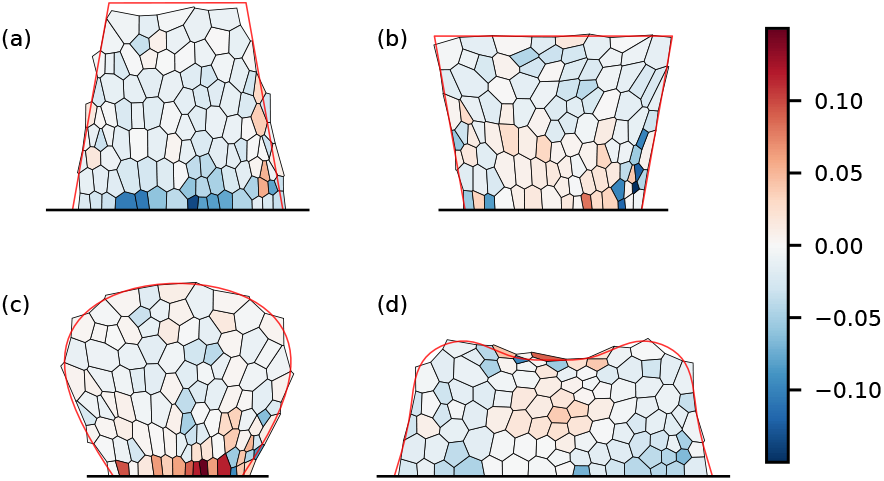
Optimization for different target boundaries (in red). The colors indicate the logarithmic deviations in measured areas compared to the goal areas. **(a)** Tall isosceles trapezoid. **(b)** Wide isosceles trapezoid. **(c)** Petal. **(d)** Non-convex shape.

As discussed, the mapping from the `MORPH` program to the target boundary is many-to-one. This allows a large range of possible distributions of polygon sizes and shapes in the model output. Although the choice of initial parameters will bias the output distribution, this choice could also be seen as a way to intentionally constrain the output space to solutions that align better with biological reasoning and experimental data. For example, the local distributions of cell areas and shape anisotropies in real plant cells are usually very narrow (barring specialized scattered cell types like stomata or the giant cells in sepals [24]). For most lateral organs, these distributions should also be symmetric with respect to the proximal-distal axis. Additionally, tissues are often divided into two or more regions in which cells have distinct identities, with size and shape characteristics that differ categorically between the regions [25, 26]. To constrain the solution space of the shape optimization to more biologically realistic configurations, we now utilize this knowledge and introduce additional (soft) constraints.

### Constraining the model output space

Thus far, we have considered a single morphogenetic target, namely, the tissue shape. However, in reality, multiple selection pressures often act simultaneously and interdependently. Our framework does accommodate this, as any number of differentiable loss terms can be included. Here, we consider including polygon metrics as a second target to be encoded in the loss function. First, we introduce a simple bias to lower the variance of the areas of the polygons in the entire tissue. Then, we further divide the tissue into regions, and investigate the effects of biasing the output separately for each region, mimicking distinct cell-type behavior.

#### Area variance

As a starting point, we consider a simple penalty on the variance of the measured polygon areas. We add a term to the loss in Eq. 7, and subsequently compute the total loss as 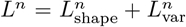, where *L*_var_ = *w*_var_ *·* Var(*a*_*i*_). Here, *w*_var_ is a parameter that controls the weight of the loss on the variance. Fig. 4 shows the effects of varying this parameter. Increasing *w*_var_ lowers overall variance, whereas the loss on the shape optimization increases on average. As expected, there is a clear trade-off between constraining the variance on the areas and the degree to which the resulting tissue shape matches the target shape. However, it is still possible to find acceptable solutions, where the optimized meshes both overlap with the target shape and have lower polygon variation. The inset figures show this qualitatively. The figure corresponding to *w*_var_ = 0 shows a high degree of overlap between the mesh and the target boundary, although with high local variation in polygon areas. At *w*_var_ = 1 the variance has been significantly reduced, but the overlap between the mesh and the target is slightly lower. Lastly, for *w*_var_ = 5, the variance is reduced further, but the shape of the mesh now differs substantially from the target boundary.

**FIG. 4:**
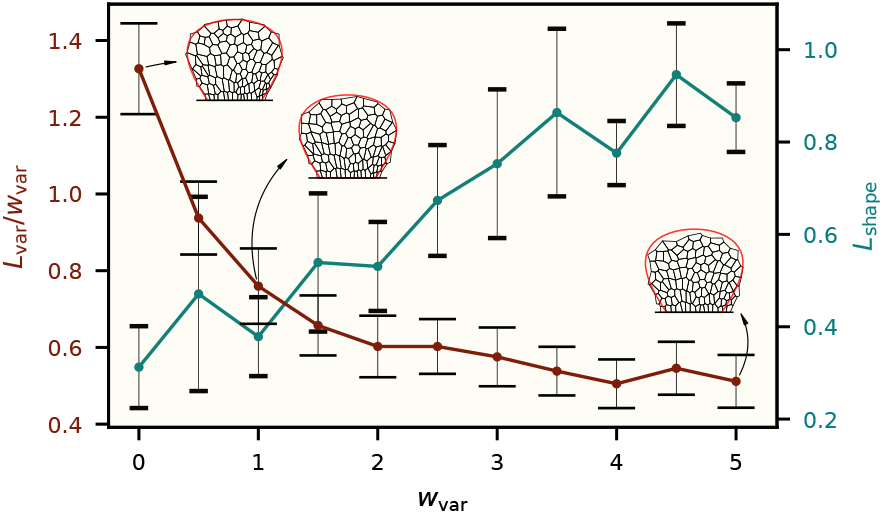
Variance in polygon areas and *L*_shape_ as a function of the parameter *w*_var_, averaged over 10 different tissue samples. Increasing the weight of the variance loss term lowers overall area variance, but increases the loss on the shape.

#### Regional cell identities

Next, we divide the tissue into multiple regions to which we assign distinct identities. The identities are characterized by biases imposed on the values of the polygon areas or anisotropies, e.g., inducing the polygons within one specific region to be particularly oblong or acquire a specific mean value of areas. We then run the shape optimization as usual with an auxiliary loss term to that of Eq. 7, to give a total loss 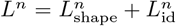. Here, *L*_id_ encodes the biases from a specific cell type configuration in a tissue, and will be defined explicitly in each of the following two examples.

In the first example, we take inspiration from the transition stage of the *Arabidopsis thaliana* embryo, where cells located close to the apico-basal axis are significantly more oblong in *y* than are the surrounding cells [27]. We thus designate the polygons as either *mid* or *outer* based on the distance in *x* between the centroids of the initial mesh and the apico-basal axis, with the cutoff set to *d*_mid_. To mimic the oblong cell type, we define a simple loss term

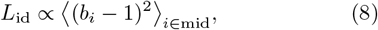

which induces the *mid* polygons to become highly *y*-anisotropic. Fig. 5a shows the measured anisotropies for the resulting mesh for the mid/outer system. First, we note that the mesh boundary largely overlaps with the target boundary, so this is a valid solution to the shape optimization problem. Second, we observe that the middle polygons are significantly narrower than those of the unbiased system (Fig. 5c), whereas the outer polygons are wider. Fig. 5b (d) shows the difference between the values in Fig. 5a (c) and those of the unbiased system in Fig. 5c, and thus demonstrates the effect of the added loss term *L*_id_. Here, we see abrupt transitions at the boundaries between the two regions, and that the bias has a greater effect for the *mid* type cells in the upper half of the mesh, where the polygons in the unbiased system are wider on average to accommodate the shape of the petal.

**FIG. 5:**
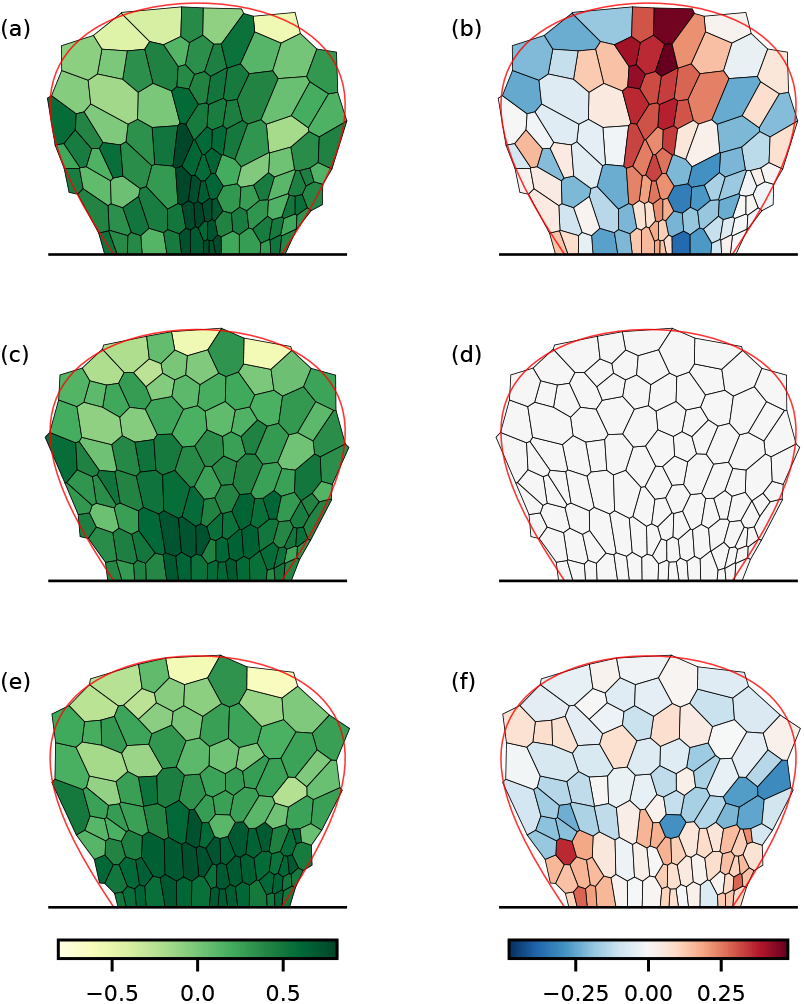
Measured anisotropies and their differences from the unbiased system for **(a-b)** the mid/outer system (*d*_mid_ = 3.0), **(c-d)** the unbiased system, and **(e-f)** the proximal/distal system (*d*_prox_ = 4.0).

In the second example we again divide the mesh into two distinct patches, this time representing tissues where the cells closest to the stem are significantly longer than the distal cells [28]. We designate the polygons as either *proximal* or *distal* based on the distance in *y* between the centroids of the initial mesh and the base, with the cutoff set to *d*_prox_. Analogous to the previous example, we add to the loss in Eq. 7 the auxiliary term

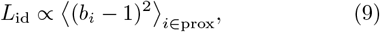

which should yield a higher anisotropy on average for the proximal polygons. The output for this system can be seen in Fig. 5e. The solution again matches the target boundary shape well, which indicates that the solution is valid for the shape optimization problem. As in the first example, we also observe two clearly defined regions, each with characteristic anisotropies. Here, the proximal cells are significantly more oblong than the distal cells, and we see abrupt changes at the border between the two regions. Fig. 5f shows the difference between the values in the proximal/distal system and the unbiased system. Compared to those of the unbiased system, the *proximal* type cells are on average more *y*-anisotropic, although half the cells do not differ significantly. This is most likely because the cells also need to accommodate the shape loss term, and are thus already more oblong in *y*.

In summary, these examples show that it is possible to train the model through direct gradient descent to both match a predetermined shape and yield a distribution of polygons that satisfies simple biological constraints.

## DISCUSSION

Here we have proposed a top-down approach to plant tissue growth that produces a final shape, using a vertex model. We applied differentiable programming to learn the parameters, and we used a minimal model, restricting the parameters for each polygon (cell or collection of cells) to one area and one shape parameter. Notably, the parameter space is large, and the parameters are thus difficult to learn because of the high non-convexity of the problem. Nevertheless, differentiable programming accelerates the search by providing gradients through the full simulation. Our exploration suggests that differentiability is most effective here as a means of fine-tuning vertex-model parameters from reasonable initial guesses, while large shape transformations remain difficult to discover from arbitrary starting points. We approached the problem of high model output degeneracy by adding an appropriate term to the shape loss function (Eq. 7). First, we restricted the amount of variance in the polygon areas across the tissue, and second, we constrained parameter properties within predefined regions in the tissue. The second approach is appropriate when some information on regional identity is already known, or can be a useful way to test hypotheses about such regional identities. Both approaches phenomenologically model the local coordination of cell behavior in biological tissues occurring through chemical and mechanical signaling.

One important factor in tissue growth is cell division: the material properties and orientation of the new cell wall that separates the two daughter cells can significantly affect the material properties of the tissue [29, 30]. However, in regions where cells have differentiated, growth mainly occurs due to cell expansion, caused by cell wall loosening and turgor pressure [31]. The distribution of dividing and growing cells in the tissue is thus time-dependent and complex. Presently, the `MORPH` program only allows for a restricted class of transformations due to inherent model limitations, resulting in polygons that are predominantly convex (see Fig. 3), and fixed in number, as we do not employ remeshing. This is particularly evident when the difference between the shapes of the initial mesh and the target boundary is too significant. One possible direction for future work is therefore to incorporate explicit cell division and growth via an iterative two-step procedure in which differentiable programming optimizes a fixed division program, over-sized cells are assigned division times, and the updated program is re-optimized.

There are a number of other logical next steps that could be taken. First, we expect that the method developed here will be particularly useful for predicting tissue growth when supplemented by experimental data. One example is the analyses of cell lineages – clonal sectors – in tissues, that display the actual patterns of growth. Differences between the model prediction and the experimental data will reveal additional constraints that can arise from the particular developmental mechanism at play, and that can subsequently be tested experimentally. Second, we envision that this method will provide additional insight when used together with models that explicitly simulate the evolution of genetic mechanisms for tissue development. Evolutionary processes do not necessarily optimize towards the most elegant mechanism for generating adaptive shapes, but are constrained by their evolutionary history (what developmental processes are already present) and by requirements for developmental and mutational robustness. Differences between predictions of our model and those of evolutionary algorithms could therefore point to the existence of such additional demands [32]. Third, to make the model generalizable and scalable, the parameters of each individual polygon could be replaced by those associated with a smaller set of points, and the polygon metrics could be calculated from these based on their positions relative to these points. The smaller parameter set would effectively act as knots in a tissue-wide function, analogous to the concentration fields of signaling molecules. This approach has several advantages, the most obvious being the fact that the total number of parameters can be kept fairly low, and independent of the polygon density. The knots could be further replaced by full (differentiable) models of gene-regulatory networks of morphogens controlling the same tissue-wide behavior. In that case, differences between the hypothesis-free differentiable model and the forward-evolution model will point to additional constraints due to the connection between pattern and growth evolution.

This work was supported by the Novo Nordisk Foundation under Grant Agreement NNF20OC0062047.

## References

[1] S. Alt, P. Ganguly, and G. Salbreux, Philosophical Transactions of the Royal Society B: Biological Sciences 372, 20150520 (2017).

[2] A. G. Fletcher, M. Osterfield, R. E. Baker, and S. Y. Shvartsman, Biophysical Journal 106, 2291 (2014).

[3] M. Marconi and K. Wabnik, Frontiers in Plant Science 12, 10.3389/fpls.2021.746183 (2021).

[4] R. Farhadifar, J.-C. Röper, B. Aigouy, S. Eaton, and F. Jülicher, Current Biology 17, 2095 (2007).

[5] K. Alim, O. Hamant, and A. Boudaoud, Frontiers in Plant Science 3, 10.3389/fpls.2012.00174 (2012).

[6] R. J. Dyson, G. Vizcay-Barrena, L. R. Band, A. N. Fernandes, A. P. French, J. A. Fozard, T. C. Hodgman, K. Kenobi, T. P. Pridmore, M. Stout, D. M. Wells, M. H. Wilson, M. J. Bennett, and O. E. Jensen, New Phytologist 202, 1212 (2014).

[7] F. Boudon, J. Chopard, O. Ali, B. Gilles, O. Hamant, A. Boudaoud, J. Traas, and C. Godin, PLOS Computational Biology 11, e1003950 (2015).

[8] I. Kneuper, W. Teale, J. E. Dawson, R. Tsugeki, E. Katifori, K. Palme, and F. A. Ditengou, Journal of Experimental Botany 72, 1151 (2021).

[9] R. M. H. Merks, M. Guravage, D. Inzé, and G. T. S. Beemster, Plant Physiology 155, 656 (2011).

[10] I. Cheddadi, M. Génard, N. Bertin, and C. Godin, PLOS Computational Biology 15, e1007121 (2019).

[11] A. Singh, A. Krishna, A. Amiri, A. Materne, P. Incardona, C. Duclut, C. M. Duque, A. Szalapak, M. Bahadorian, S. K. T. Veettil, et al., arXiv preprint 2509.24905 (2025).

[12] M. Marin-Riera, M. Brun-Usan, R. Zimm, T. Välikangas, and I. Salazar-Ciudad, Bioinformatics 32, 219 (2016), https://academic.oup.com/bioinformatics/article-pdf/32/2/219/49016459/bioinformatics322219.pdf.

[13] M. H. Swat, G. L. Thomas, J. M. Belmonte, A. Shirinifard, D. Hmeljak, and J. A. Glazier, in Computational Methods in Cell Biology, Methods in Cell Biology, Vol. 110, edited by A. R. Asthagiri and A. P. Arkin (Academic Press, 2012) pp. 325–366.

[14] S. Green and R. Batterman, Studies in History and Philosophy of Science Part C: Studies in History and Philosophy of Biological and Biomedical Sciences 61, 20 (2017).

[15] A. G. Fletcher and J. M. Osborne, WIREs mechanisms of disease 14, e1527 (2022).

[16] E. Coen and D. J. Cosgrove, Science 379, eade8055 (2023).

[17] T. Lecuit and L. Le Goff, Nature 450, 189 (2007).

[18] G. A. Stooke-Vaughan and O. Campàs, Current opinion in genetics & development 51, 111 (2018).

[19] L. Happel and A. Voigt, Journal of Nonlinear Science 36, 15 (2026).

[20] I. Burda, A. C. Martin, A. H. Roeder, and M. A. Collins, Developmental cell 58, 2850 (2023).

[21] R. Deshpande, F. Mottes, A.-D. Vlad, M. P. Brenner, and A. Dal Co, Nature Computational Science 5, 875 (2025).

[22] J. Bradbury, R. Frostig, P. Hawkins, M. J. Johnson, C. Leary, D. Maclaurin, G. Necula, A. Paszke, J. VanderPlas, S. Wanderman-Milne, and Q. Zhang, jax: composable transformations of python+numpy programs (2018).

[23] H. Ronellenfitsch and E. Katifori, Physical review letters 117, 138301 (2016).

[24] A. H. K. Roeder, Quantitative Plant Biology 2, e14 (2021).

[25] E. Salvi and E. Moyroud, The Plant Journal 121, e70101 (2025), eprint: https://onlinelibrary.wiley.com/doi/pdf/10.1111/tpj.70101.

[26] E. Doody and E. Moyroud, New Phytologist 247, 2538 (2025), eprint: https://nph.onlinelibrary.wiley.com/doi/pdf/10.1111/nph.70370.

[27] C. A. ten Hove, K.-J. Lu, and D. Weijers, Development 142, 420 (2015), https://journals.biologists.com/dev/article-pdf/142/3/420/1840396/dev111500.pdf.

[28] L. Riglet, A. Zardilis, A. L. Fairnie, M. T. Yeo, H. Jönsson, and E. Moyroud, Sci Adv 10, eadp5574 (2024).

[29] P. Sahlin, O. Hamant, and H. Jönsson, in Complex Sciences, edited by J. Zhou (Springer Berlin Heidelberg, Berlin, Heidelberg, 2009) pp. 971–979.

[30] A. Bonfanti, E. T. Smithers, M. Bourdon, A. Guyon, P. Carella, R. Carter, R. Wightman, S. Schornack, H. Jönsson, and S. Robinson, Proceedings of the National Academy of Sciences 120, e2302985120 (2023), https://www.pnas.org/doi/pdf/10.1073/pnas.2302985120.

[31] S. McQueen-Mason, D. M. Durachko, and D. J. Cosgrove, The Plant Cell 4, 1425 (1992), https://academic.oup.com/plcell/article-pdf/4/11/1425/34745719/plcellv4111425.pdf.

[32] R. M. A. Vroomans, P. Hogeweg, and K. H. W. J. ten Tusscher, EvoDevo 7, 14 (2016).

