## Supplementary Material for "Differentiable Vertex Model: Exploring Gradient-Based Optimization for Tissue Morphogenesis"

Lars Erik J. Skjegstad
*Niels Bohr Institute, University of Copenhagen, Denmark*

Steven Oud and Renske M. A. Vroomans
*Sainsbury Laboratory, University of Cambridge, U.K.*

Julius B. Kirkegaard*
*Niels Bohr Institute, University of Copenhagen, Denmark and*
*Department of Computer Science, University of Copenhagen, Denmark*
(Dated: May 1, 2026)

### CONTENTS

I. Source code 2

II. Optimization pipeline 2

III. Shape optimization details 3
A. Optimization procedure 3
B. Metrics and residual discrepancies 3

IV. Model limitations 4

References 4

### I. SOURCE CODE

The source code used for all simulations is available at `https://github.com/larserik-js/diff-tissue`. The version used for this paper corresponds to the Git tag `v1.5.0`.

### II. OPTIMIZATION PIPELINE

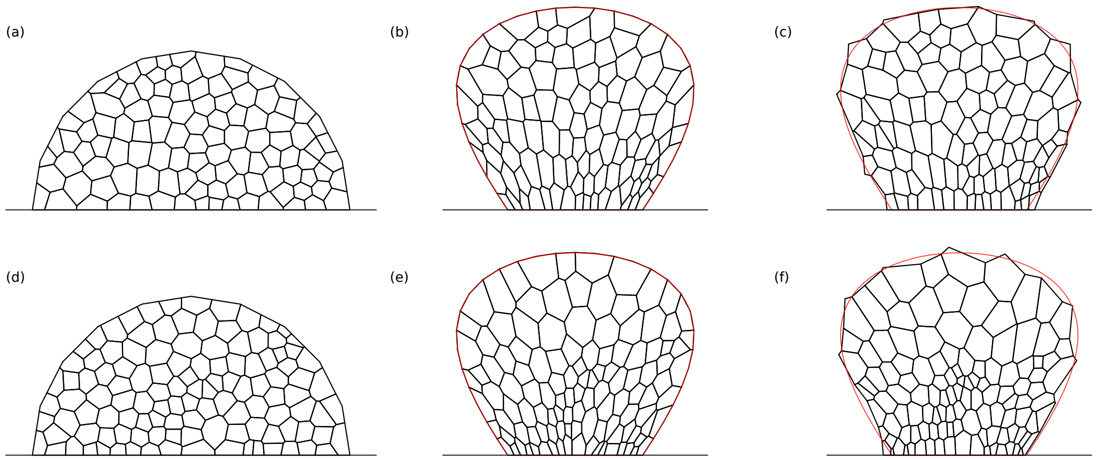


FIG. S1. Optimization pipeline. **(a)** Initial mesh, seed = 0. **(b)** Tutte embedding of the initial mesh in (a). **(c)** Optimized mesh, unbiased system. **(d)** Initial mesh, seed = 1. **(e)** Tutte embedding of the initial mesh in (d). **(f)** Optimized mesh, proximal/distal system.

Fig. S1 shows the optimization pipeline for two different initial mesh initializations. We embed the initial mesh (Fig. S1a) onto the target boundary using the Tutte embedding (Fig. S1b). We then use the measured metrics from the Tutte embedding as the initial parameters for the shape optimization, the output of which can be seen in Fig. S1c. Figs. S1d-f show the analogous plots where the output distribution is biased toward the proximal/distal configuration during optimization. The target shape is shown in red.

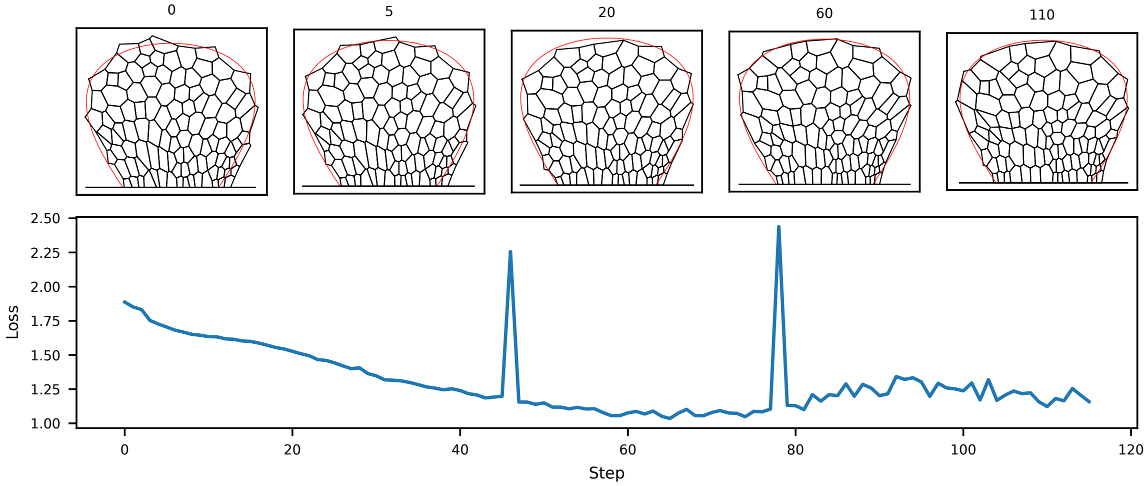


FIG. S2. Different steps of the optimization procedure for the proximal/distal cell configuration, as well as the corresponding loss values.

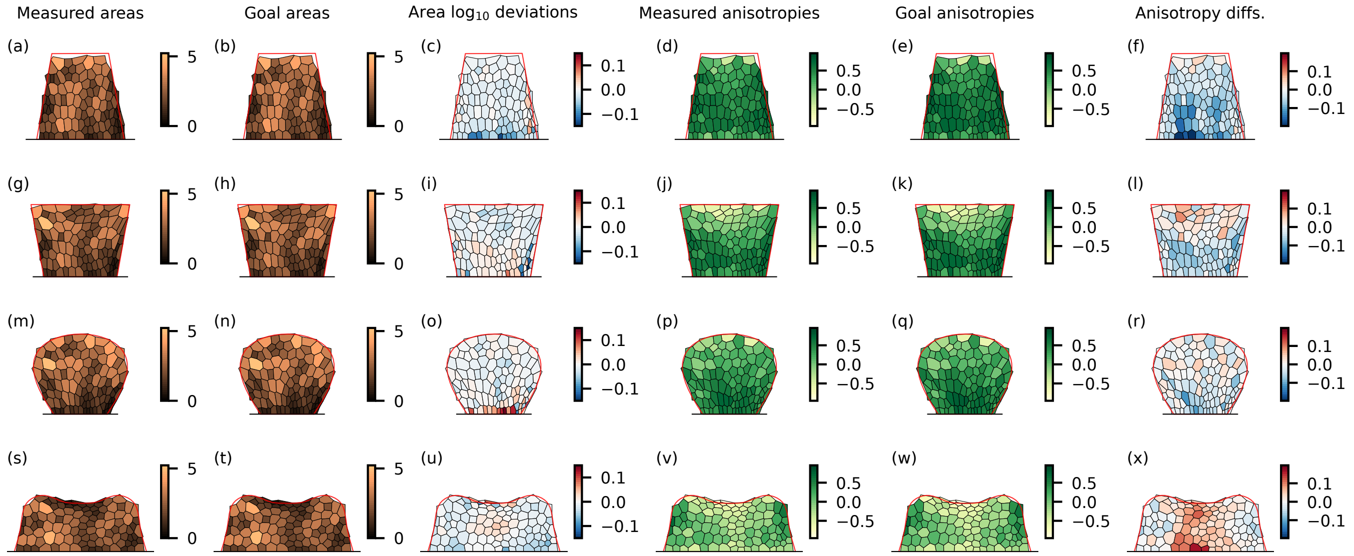


FIG. S3. Measured/goal areas and anisotropies and their deviations for four different target shapes.

### III. SHAPE OPTIMIZATION DETAILS

#### A. Optimization procedure

During the optimization, we transform the areas and anisotropies to logits, the domain of which is the entire real number line. The values are converted to goal metrics by using sigmoids (and their inverse when converting back to logits), where the minimum and maximum values constitute parameter constraints. The goal areas are limited to the range $[0, 5.0 \cdot \max(a_i^{t=0})]$, whereas anisotropies are constrained to the values $[-1, 1]$ by definition.

We stop the optimization procedure after 50 steps of non-decreasing loss occur, and keep the solution with the lowest loss value.

Fig. S2 shows different steps of the optimization procedure for the proximal/distal cell configuration. The first figure (step number 0) shows the results after 0 parameter updates, i.e., using the inputs from the Tutte embedding. The bottom figure shows the loss value as a function of iteration steps.

#### B. Metrics and residual discrepancies

Fig. S3 shows a detailed plot of the values for the optimized mesh. The figure compares measured and goal values for both areas and anisotropies, for four different target boundaries. In addition, the residual discrepancies for both metrics can also be seen.

### IV. MODEL LIMITATIONS

The model accuracy depends heavily on the target shape in question. To quantify this, we define an isosceles trapezoid as target shape, where the counterclockwise angle between the base and the right leg is an input parameter. We then run 10 simulations for each target trapezoid, and compute the mean loss. Fig. S4 shows the results. As expected, the results are significantly worse for the triangle shape (angle = 120 degrees) and upside-down triangle shape (angle $\rightarrow$ 60 degrees). This indicates that there are limits to the types of target shapes to which the initial mesh can transform for a fixed topology.

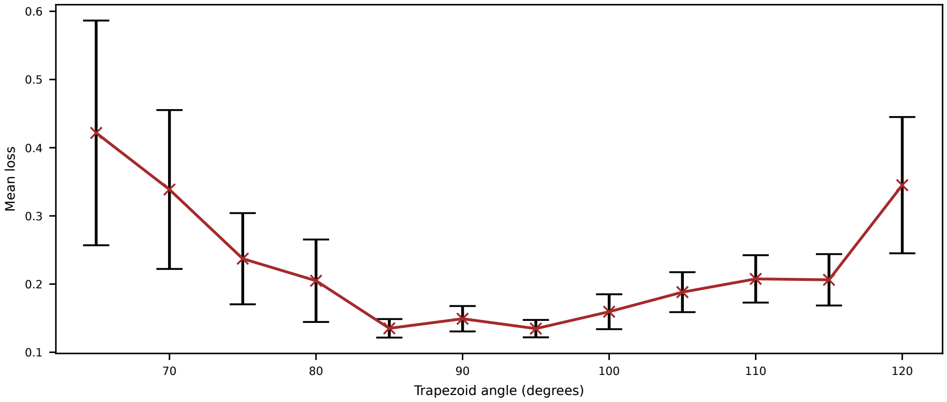


FIG. S4. Mean loss with standard errors as a function of angle for the parameterized isosceles trapezoid. The values are calculated on the results from 10 different random initial meshes.
